# The cryo-EM structure of human CST reveals a two-megadalton decameric assembly bound to telomeric DNA

**DOI:** 10.1101/815886

**Authors:** Ci Ji Lim, Alexandra T. Barbour, Arthur J. Zaug, Allison E. McKay, Deborah S. Wuttke, Thomas R. Cech

**Affiliations:** Department of Biochemistry, University of Colorado Boulder, CO 80303, USA; BioFrontiers Institute, University of Colorado Boulder, CO 80303, USA; Howard Hughes Medical Institute, University of Colorado Boulder, CO 80303, USA

## Abstract

The single-stranded DNA-binding CTC1-STN1-TEN1 (CST) complex is essential for telomere maintenance and genome-wide replication recovery, processes that are critical for genome stability. Here, we report the 2.95 Å cryo-EM structure of human CST bound to telomeric single-stranded DNA, which unexpectedly assembles as a decameric supercomplex. The atomic model of the 134 kDa CTC1, built almost entirely *de novo*, reveals the overall architecture of CST and the DNA-binding anchor site. In situ arrangements of STN1 and TEN1 are revealed, with STN1 interacting with CTC1 at two separated sites, allowing allosteric mediation of CST decameric assembly. Surprisingly, CTC1 lacks the anticipated structural homology to yeast Cdc13 but instead shares similarity with a form of Replication Protein A. The atomic-resolution model of human CST provides crucial mechanistic understanding of CST mutations associated with human diseases. Moreover, the decameric form of CST suggests the intriguing possibility of ssDNA architectural organization similar to what the nucleosome provides for dsDNA.

**One Sentence Summary:** Human telomeric single-stranded DNA triggers the assembly of a decameric protein supercomplex solved by cryo-EM.

---

CST is a heterotrimeric ssDNA-binding complex present in many species (*1*–*3*). Although CST was initially discovered as a DNA polymerase-primase cofactor (*4*) and telomere-associated protein complex (*2*, *5*–*7*) essential for telomere replication, its role has now expanded genome-wide, with CST key in recovering stalled replication forks (*2*, *8*–*11*) and facilitating DNA damage repair (*12*–*15*). Consequently, mutations in CST are the basis of human genetic diseases such as Coats Plus Syndrome and Dyskeratosis congenita (*16*–*21*).

Although CST is similar to the telomere-associated POT1 regarding preferential binding of G-rich ssDNA (*22*–*25*), CST is also able to bind non-specific sequences within longer ssDNA (*6*, *25*), analogous to human RPA. Studies have tried to determine the DNA-binding region(s) of CST to shed light on its mechanistic functions but have been hampered by poor understanding of mammalian CST architecture, especially because an intact heterotrimeric complex is necessary for its DNA-binding function (*6*, *17*, *26*). Crystal structures of human components are limited to STN1 and TEN1 (*26*), while the structure of the largest subunit CTC1 has proven technically challenging to solve, and only a single OB domain has been determined (*27*). The yeast Cdc13 protein also associates with Stn1 and Ten1 and has therefore been proposed as a CTC1 homolog, but Cdc13 and mammalian CTC1 are unrelated in sequence. Hence, it is unclear if Cdc13 and CTC1 share structural homology. It is, therefore, critical to determine the atomic-resolution architecture of mammalian CST, especially given its expanding roles in telomere and genome-wide DNA replication and repair pathways.

## Cryo-EM structure of human CST decameric supercomplex

We solved the structure of purified recombinant human CST protein (hereafter termed DNA-free CST) to 7 Å resolution using single-particle cryo-EM (fig. S1). To improve the resolution of the structure, we added a minimal telomeric ssDNA (3xTEL, (TTAGGG)_3_) (*6*) to the purified CST protein and unexpectedly discovered a symmetric complex dramatically larger than the monomeric CST during cryo-EM analysis. The supercomplex was also observed by size exclusion chromatography (**Fig. 1A**).

**Fig. 1.**
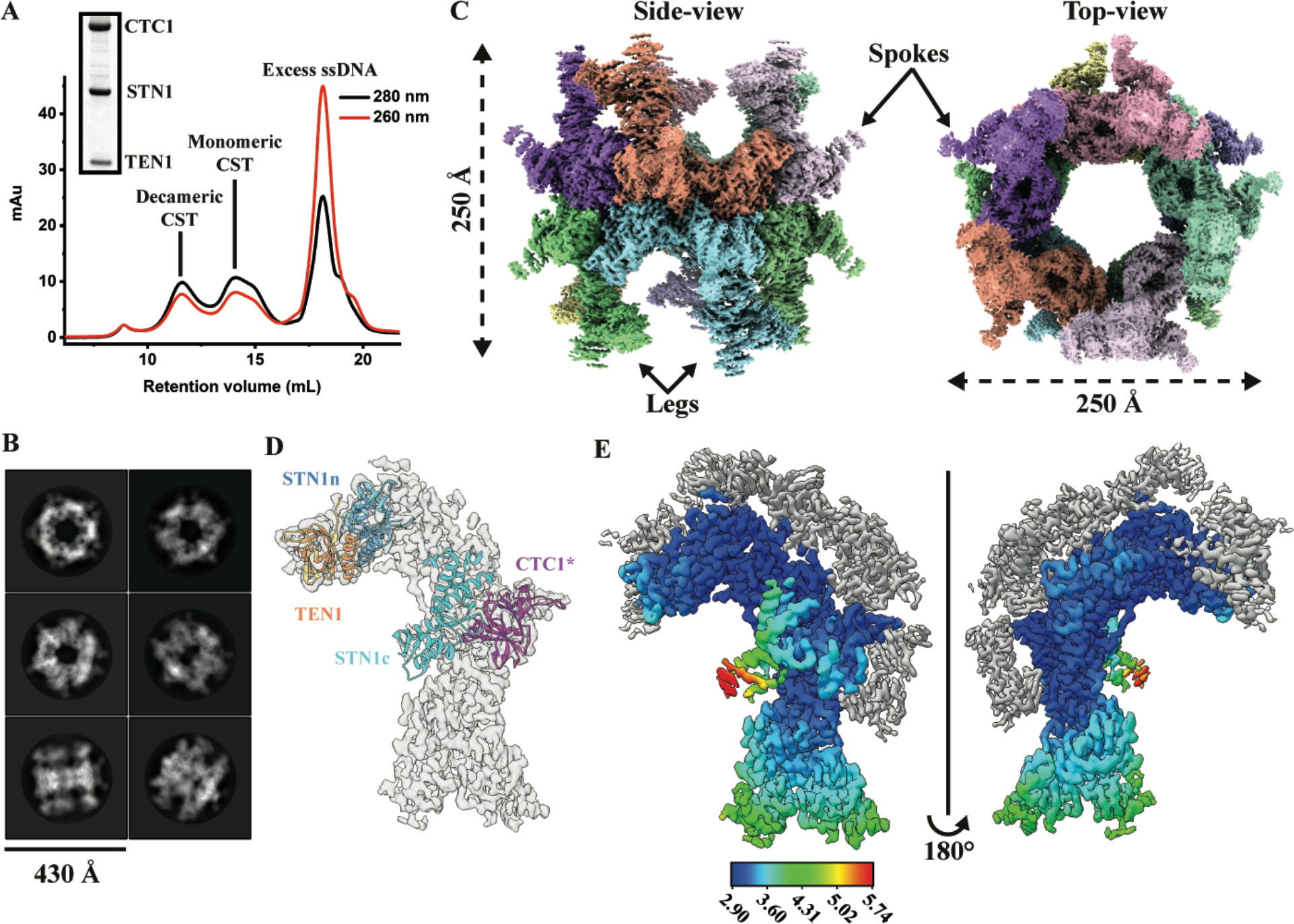
Cryo-EM structure of human CST decameric supercomplex. **(A)** Size-exclusion chromatography of human CST-ssDNA complexes resolves various oligomeric states. Inset, SDS-PAGE Coomassie-stained bands of the purified CST heterotrimeric complex – CTC1 (~135 kDa), STN1 (~42 kDa) and TEN1 (~14 kDa) **(B)** Representative two-dimensional (2D) classes of CST-3xTEL particles. **(C)** Cryo-EM density of decameric CST complex colored by segmented CST monomers. **(D)** Docking of available atomic models of a CTC1 OB-domain (* reported as central domain OB-fold (*27*), PDB 5W2L), STN1n (N-terminal half, PDB 4JOI:A), STN1c (C-terminal half, PDB 4JQF) and TEN1 (PDB 4JOI:C). **(E)** Cryo-EM density of refined CST monomer after symmetry expansion and three-dimensional (3D) classification. The grey density belongs to neighboring CST monomers in the decameric supercomplex. The model is colored based on local resolution (rainbow color scale-bar).

We collected two cryo-EM movie datasets of the CST-3xTEL sample at 0° and 30° stage-tilts to resolve preferred-orientation problems of vitrified particles (*28*) (fig. S2). This led to a 3D reconstruction of the decameric CST supercomplex at 3.03 Å global resolution (**Fig. 1B and C**, fig. S3). D5 symmetry was imposed during 3D refinement, which improved the resolution but could average-out different monomeric CST conformations in the decameric supercomplex (*29*). Therefore, we performed D5 symmetry expansion before using 3D classifications with a monomeric CST mask (derived from our DNA-free CST model, fig. S1) to tease out a more homogenous population of CST monomers. The selected intact CST monomeric particles were then subjected to 3D refinement (now with no symmetry imposed, using C1 symmetry) with a mask that was expanded to include the monomer and surrounding interfaces with adjacent monomers. (fig. S4). Although there was a slight decrease in map resolution to 3.06 Å, the quality of the electron density map was significantly improved visually (fig. S5A).

As evident from the 2D classification and the consensus decameric model (**Fig. 1B & C**), the spokes and legs of the decameric CST complex are highly flexible, which translates to poorer resolution in the consensus model map (fig. S5B). Still, the high-resolution EM map allowed docking of the crystal structures solved for domains of human TEN1, STN1 (*26*) and a central OB-domain (oligonucleotide/oligosaccharide-binding fold) of CTC1 (*27*) with high confidence (**Fig. 1D**). Because the spoke density was accounted for by the docking of the crystal structure of STN1 (*26*), we focused on improving the resolution of the leg density (fig. S4). Focused 3D classifications on the legs yielded a single 3D class (~50% population) that presumably had a well-defined legs conformation, and masked 3D refinement (using the entire monomer mask) on this subset of the particles resulted in a 2.95 Å global resolution map (**Fig 1E**) with local resolution of the majority of the legs significantly improved to ~4.5 Å.

## Overall architecture of human CST

The symmetry-expanded CST electron density map also enabled *de novo* building of the entire CTC1 atomic model (894 residues, excluding the one CTC1 OB-domain solved (*27*)), revealing the overall architecture of the human CST complex (**Fig. 2A**, fig. S5C, table S1). CTC1 is comprised of an unprecedented 7 tandem OB-domains (OB-A through G, **Fig. 2B & C**). The intricacy of the CST structure explains why full-length CTC1 is critical for CST assembly and ssDNA binding, as well as why isolating individual predicted CTC1 OB-domains for structural studies has been technically challenging (*2*, *26*, *27*). The human CST complex has a subunit stoichiometry of 1:1:1, unlike that reported for *Candida glabrata* CST complex (*30*).

**Fig. 2.**
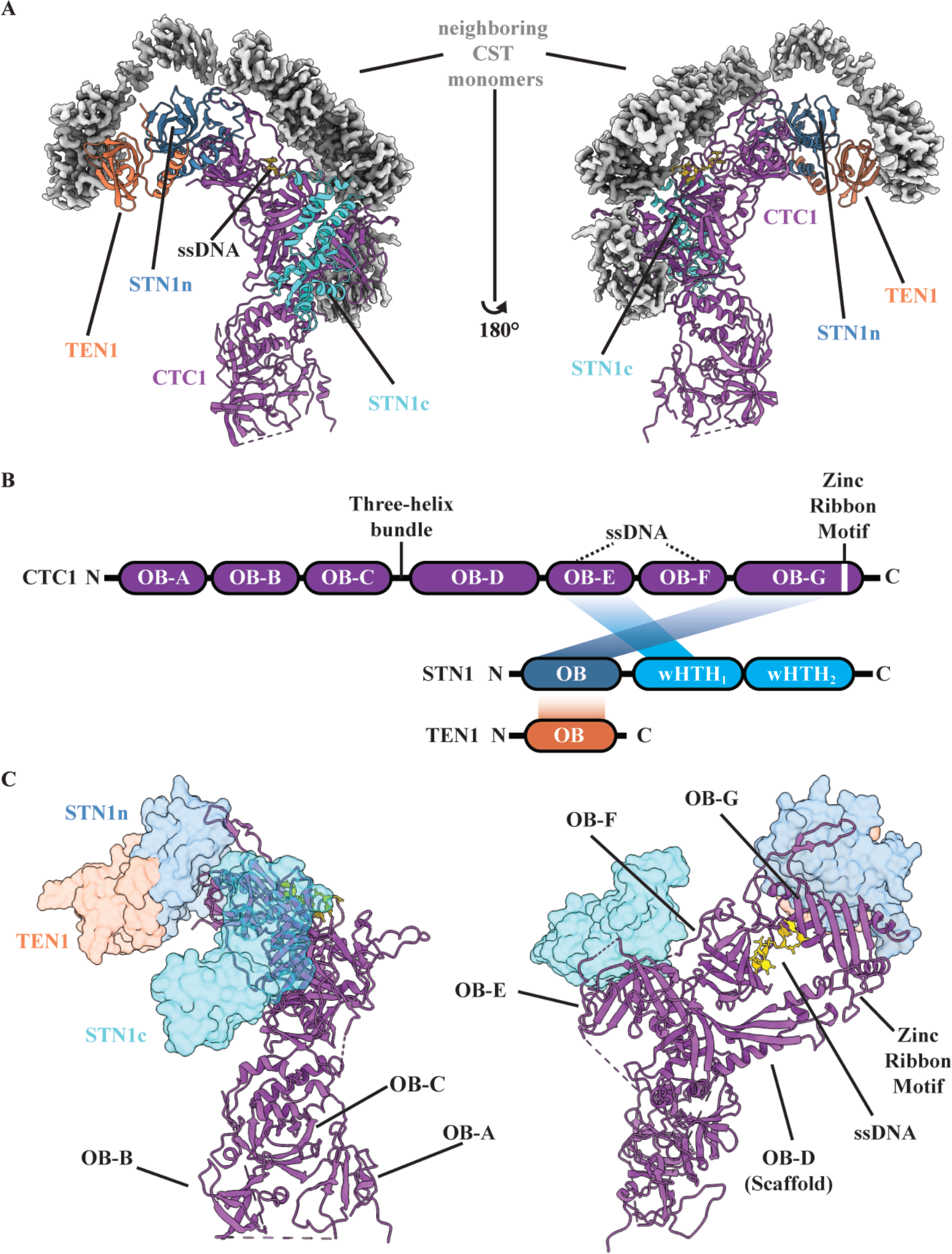
Architecture of human CST and atomic model. **(A)** Atomic model of monomeric CST, with the EM density (grey) of neighboring CST molecules (partial). CST subunits are colored as – CTC1 (purple), STN1n (blue), STN1c (cyan), TEN1 (orange) and ssDNA (gold). **(B)** Structure-based schematic of CST domain architecture and inter-molecular interactions between subunits. **(C)** CTC1 architectural organiation of seven OB-domains (A-G) and identified Zinc-ribbon motif and a bound-ssDNA.

The C-terminus of CTC1 serves as a structural scaffold for STN1 and TEN1 binding (**Fig. 2A**), providing understanding of past genetic and biochemical studies (*6*, *17*). A single STN1 protein has two separate interaction sites with CTC1, with the STN1 N-terminal half (STN1n) interacting with CTC1 OB-G and the C-terminal half (STN1c) with CTC1 OB-E (**Fig. 3A**). These two halves of STN1 are connected by an unstructured peptide linker of 7 residues (**Fig. 2B**). In contrast to RPA (*31*, *32*), there is no triple-helix bundle stabilizing the heterotrimeric CST complex (**Fig. 3B**). Instead, TEN1 binding to CTC1 is bridged by STN1n (similar to the low-resolution model of the *Tetrahymena* CST (*3*, *33*)) (**Fig. 2A-C**), with STN1n binding to a highly conserved interaction patch on CTC1 OB-G (fig. S6A & B). The first winged helix-turn-helix (wHTH) domain of STN1c interacts with CTC1 OB-E (**Fig. 2B & C**), the structure of which was previously determined (*27*). However, no strong conservation of residues was found on the interaction patch of CTC1 OB-E, suggesting that STN1c-CTC1 interaction could be weaker than STN1n-CTC1 interaction, as seen with *Tetrahymena* CST complex (*33*). Supporting this hypothesis, we found STN1n alone was able to interact with CTC1, but STN1c could not (fig. S6B). In addition, TEN1 interaction with CTC1 was lost with STN1c but not with STN1n (fig. S6B), supporting the cryo-EM model in which TEN1 is bridged to CTC1 via STN1n (**Fig. 2A-C**).

**Fig. 3.**
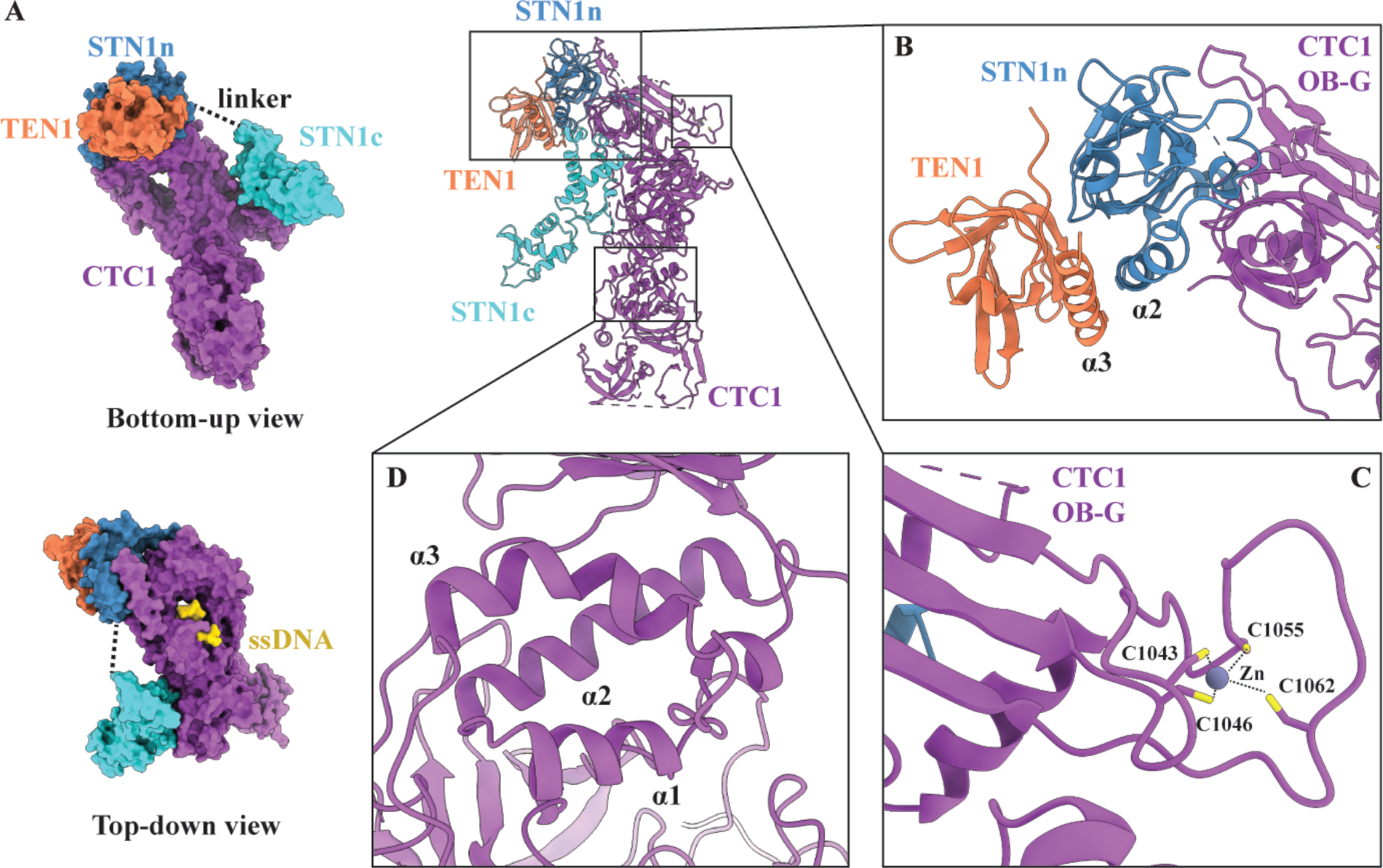
CST inter-subunit interactions and CTC1 molecular motifs. **(A)** STN1 N-terminal and C-terminal halves – STN1n and STN1c – interact separately with CTC1 via a flexible peptide linker. **(B)** STN1 and TEN1 do not interact with CTC1 using a trimeric helix-bundle like human RPA; instead, STN1 directly interacts with a highly-conserved patch on CTC1 and bridges TEN1 to CTC1. **(C)** Zinc ribbon motif within CTC1 OB-G with zinc coordinated by four conserved cysteine residues. The distances between the zinc ion and cysteines range from 2.3 to 3.6 Å. **(D)** A three-helix bundle (annotated *α*1, *α*2 & *α*3) connects OB-A, B and C to the rest of the C-terminal OB-domains of CTC1.

CTC1 OB folds E, F, and G are arranged spatially on OB-D, which acts as a scaffold, resulting in the four OBs forming a ring-like structure (**Fig. 2C**). Structural homology analysis of individual CTC1 OB domains found CTC1 to be most similar to RPA and Teb1 (RPA-like paralog in *Tetrahymena*) (fig. S7A), with CTC1 OB-F most similar to Teb1’s OB-B (*34*), and OB-G similar to OB-C of RPA70 or Teb1 (*31*). The previously solved OB-E was reported to have similarity to *Ustilago maydis* RPA70 OB-A (*27*). CTC1 OB-G also has a zinc ribbon motif like the OB-C domains of RPA and Teb1 (*31*, *34*) (**Fig. 3C**, fig S7B), suggesting CTC1 OB-G zinc ribbon motif could play a role in ssDNA-binding similar to RPA and Teb1. The scaffold OB-D has no convincing structural homology hits, but given its unique extended OB-fold structure, it could be an evolved form of the more compact and conventional OB-fold (*35*).

Given the long-standing suggestion that yeast Cdc13 and mammalian CTC1 are homologs (*36*), it is unexpected that we found no strong structural homology between them. Instead, the C-terminal half of human CST (comprised of OB-E, F and G) shares architectural similarity with RPA70. We also found a three-helix bundle bridging OB-C and OB-D (**Fig. 3D**). This three-helix bundle effectively segregates OB-A, B and C from the C-terminal OB-domains (**Fig. 2B & 3D**). For the CST “legs”, OB-C serves as a scaffold for both OB-A and OB-B (**Fig. 2B & C**). Because of extensive flexibility of OB-A, we were only able to *de novo* build a partial model for it, with the rest (~45 residues) built using as a poly-alanine model. Still, the overall model of OB-A clearly showed the OB-fold topology. Structural homology searches of OB-B and OB-C revealed them to be most similar to *Ustilago maydis* RPA70 OB-A and OB-B (*32*) (fig. S7A). The multiple structural hits of human CTC1 to various domains of RPA70 suggest CTC1 may have evolved from RPA via gene duplication.

Disease mutations in CTC1, A227V, V259M and V665G, which have been shown to interfere with pol-*α* binding (*17*), are located on CTC1 OB-B (A227 & V259) and on scaffold OB-D (V665) (fig. S8). Given that Pol-*α* has a bi-lobal architecture (*37*), the catalytic and primase lobes of Pol-*α* could engage CST at separate sites. After building the atomic-model of CTC1, we found extra density near OB-F of CTC1 that we attribute to the telomeric ssDNA (**Fig. 2B & C**). This extra density was not seen in the monomeric CST complex model obtained without DNA addition (fig. S9A).

## Anchor side of CST single-stranded DNA-binding pocket

Only four nucleotides, TAGG, are clearly visible in the EM map (**Fig. 4A-C**), which suggests that these interact with the protein most strongly or with highest occupancy, and the rest of the ssDNA is likely flexible or bound to CST in multiple binding modes. This is not unexpected given CST binds multiple configurations of ssDNA dynamically (*6*, *9*, *25*), akin to the closely related human RPA (*38*). Hereafter, we term the site of ssDNA binding on OB-F as the ssDNA anchor patch. Several positively charged residues are involved in the ssDNA anchor patch – CTC1 R978, K1164, K1167, as well as additional aromatic and neutral-polar residues – CTC1 Y949, N981, Y983 (**Fig. 4D & E**). The anchor patch utilizes several kinds of interactions between CTC1 and ssDNA, for example, Y983 hydrogen bonding to the negatively charged ssDNA phosphate backbone, N981 and K1164 hydrogen bonding to ssDNA bases, and Y949 π-π stacking with the A2 base which in turn is stacked on the T1 base (**Fig. 4E**). The tyrosine-base-base stack is reminiscent of the stacking arrangements seen in the human POT1-ssDNA structure (*22*) These residues are highly conserved across mammalian CTC1 homologs (fig. S10) and they bridge CTC1 OB-F and OB-G, suggesting that ssDNA binding can potentially stabilize CTC1 architecture (**Fig. 4D**).

**Fig. 4.**
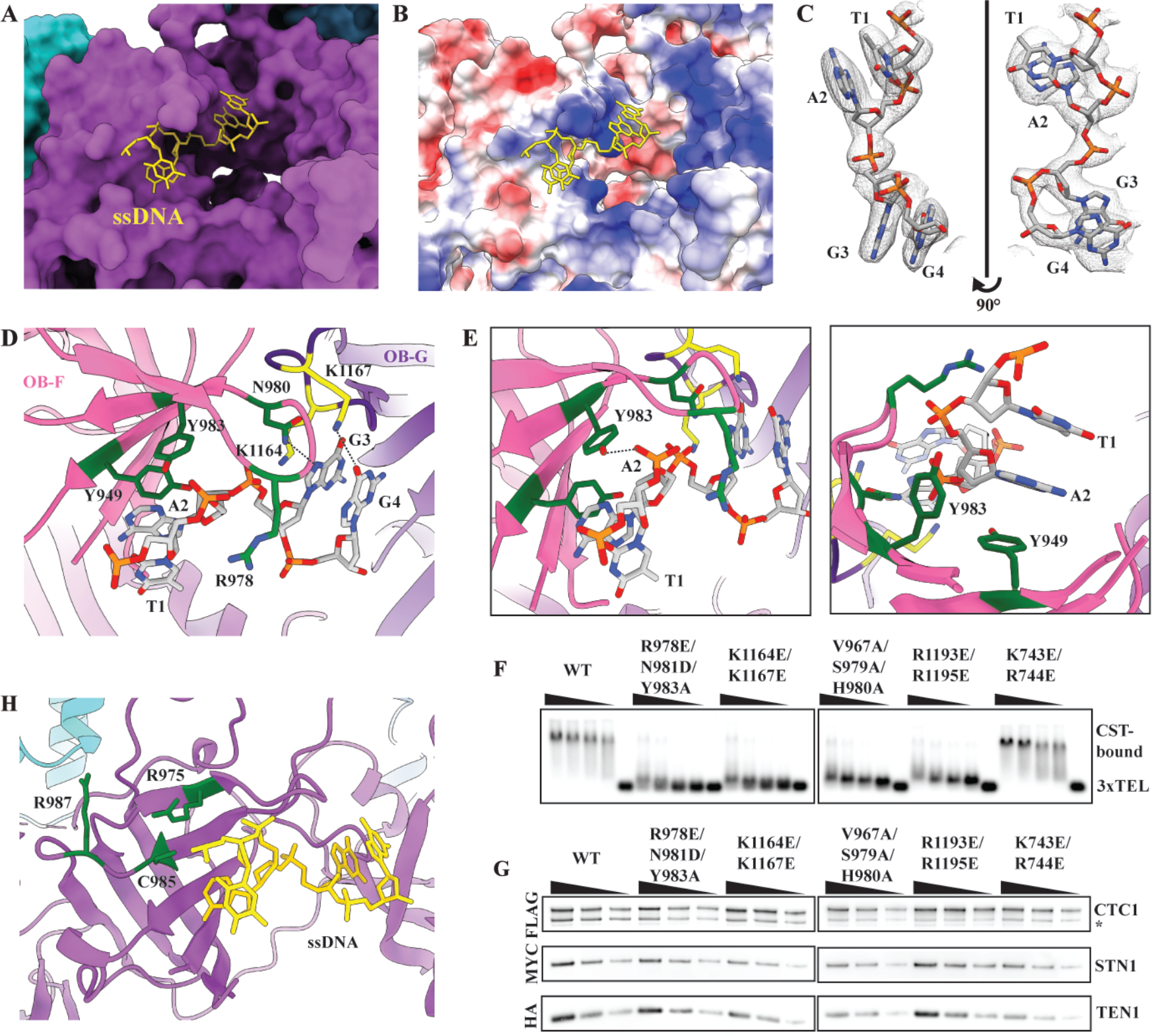
Telomeric ssDNA-binding anchor site of CST. **(A)** A 4 nt segment of the single-stranded telomeric DNA is located on CTC1. **(B)** Columbic surface analysis (*48*) reveals the ssDNA anchor site is highly positively charged (blue; red is negatively charged surface). **(C)** Cryo-EM density of the ssDNA molecule built with the sequence assigned as TAGG (5’ – T1-A2-G3-G4 – 3’). The numbering is based on the visible ssDNA, not the full-length ssDNA. **(D)** CTC1 residues involved in ssDNA-binding are shown in green and yellow from OB-F (pink) and OB-G (purple), respectively. Predicted hydrogen bonds are shown as black dashed lines. **(E)** Representative residues of CTC1 that are involved in direct interactions with the phosphodiester group of ssDNA (Y983, left panel) or π-π stacking with a ssDNA base (Y949, right panel). (**F**) Gel-shift assay showing CST DNA-binding mutants predicted from the atomic model no longer bind telomeric ssDNA (TTAGGG)_3_. **(G)** CST DNA-binding mutants are still able to form heterotrimeric CST complex as shown by tandem immunoprecipitation pull-downs (FLAG/HA) from exogenously expressed FLAG-CTC1, Myc-STN1 and HA-TEN1. * indicates protein degradation products. **(H)** Human CTC1 disease mutations (*17*) that abolish ssDNA binding (green residues) are located near the ssDNA anchor site.

To validate and determine the energetic contributions of the observed protein-DNA interactions, we performed mutagenesis on sets of CTC1 residues in the anchor patch – R987E/N981D/Y983A (anchor site on OB-F), K1164E/K1167E (anchor site on OB-G), V967A/S979A/H980A (structural integrity residues on OB-F), R1193E/R1195E (structural integrity residues on OB-G). Each of these sets of mutations abolished CTC1 DNA-binding activity, while the K743E/R744E negative control mutation did not (**Fig. 4F**, fig. S9B). Crucially, none of these sets of mutations disrupted the assembly of the heterotrimeric CST complex as shown by their co-immunoprecipitation (**Fig. 4G**). This is important given an intact complex is necessary for CST DNA binding (*39*).

Previously identified CTC1 disease mutations (R975G, C985Δ and R987W) that have been shown (*17*) to impact CST-ssDNA binding are also in the vicinity of the ssDNA anchor patch (**Fig. 4H**). An interesting hit that emerged from our structural homology search for CTC1 OB-F is *S. pombe* POT1 (*40*, *41*) (fig. S7A), which binds telomeric ssDNA at the same side of CTC1 OB-F and possesses multiple DNA-binding modes.

## Assembly mechanism and pathways of decameric CST supercomplex

The masked 3D refinement of the CST supercomplex after symmetry expansion (**Fig. 1E**) also allowed identification of interactions that appear to mediate decameric supercomplex assembly (**Fig. 4**). The sites can be categorized into two oligomerization classes (**Fig. 5B-C**): (1) dimerization (dihedral dimerization or inverted dimer) and (2) tetramerization (two sub-classes: adjacent and diagonal). For CST dimerization, three conserved residues at the interface are N745, L843 and R1175. R1175 is particularly interesting given it is within range (< 5 Å) for interaction with the phosphodiester groups of T1 or A2 of the opposite dihedral dimer’s telomeric ssDNA (**Fig. 5B**). This suggests ssDNA-binding can also stabilize CST dihedral dimerization through a charge-charge interaction. In addition, the proximity of the two ssDNA anchor patches across dihedral dimers (fig. S11) suggests the tantalizing possibility that a single molecule of ssDNA could “staple” together two dimers; consistent with this model, CST needs a minimum length of ssDNA (≥ 18 nts) for tight binding (*6*). For tetramerization, CTC1 interacts with its adjacent neighbor’s TEN1 (CTC1 H484 & R624) (**Fig. 5C**) and diagonally opposite neighbor’s STN1n (CTC1 E1183) (**Fig. 5D**), demonstrating the intricate assembly of the CST decamer.

**Fig. 5.**
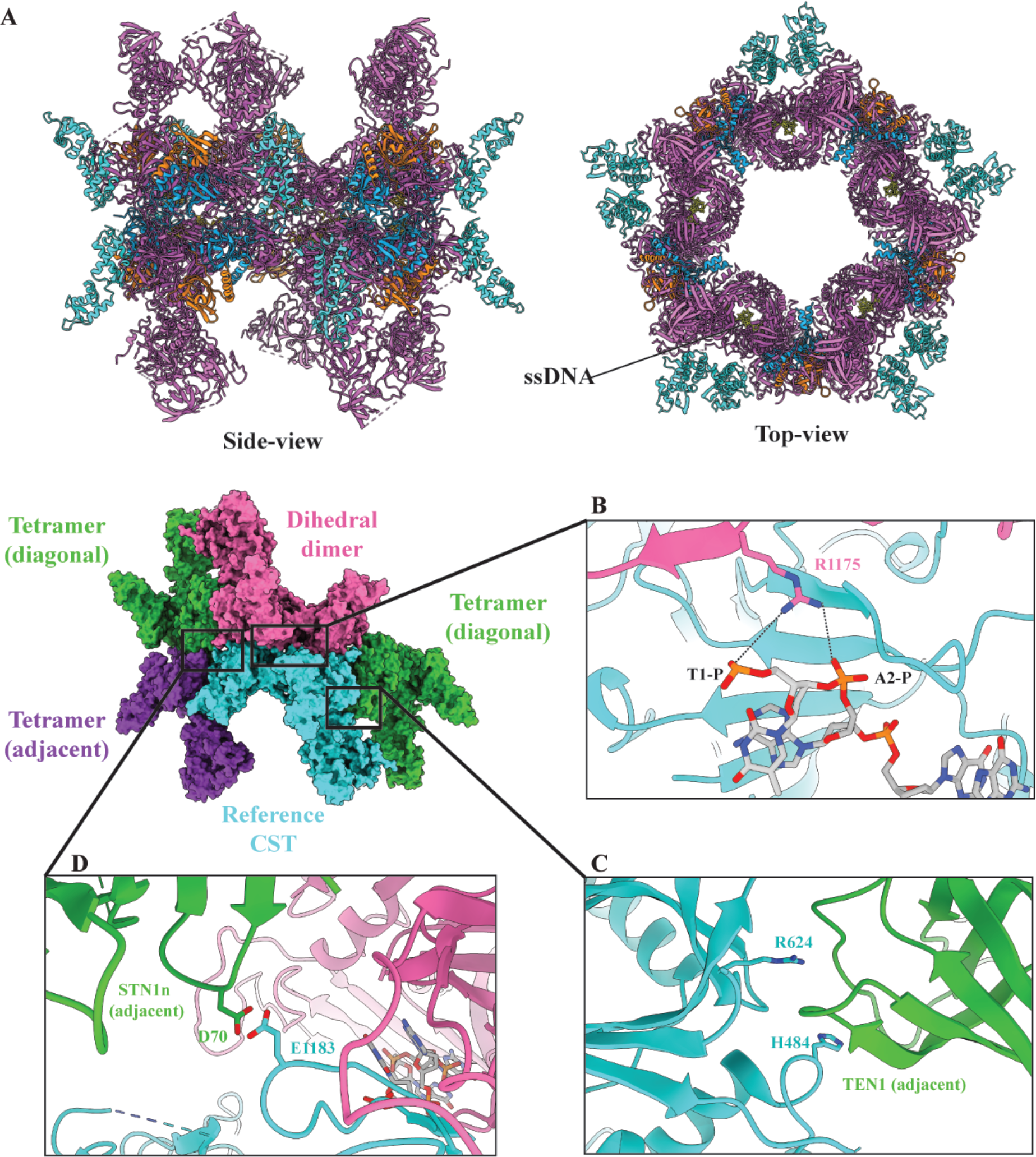
Identification of molecular interactions underlying CST decameric supercomplex formation. **(A)** Atomic models of CST arranged in decameric supercomplex. (**B-C**) The reference CST (cyan) is flanked by four CST complexes – a dihedral dimer (opposite, pink) and three tetrameric partners (two diagonal neighbors, green, and one adjacent neighbor, purple). **(B)** CTC1 R1175 from the dihedral dimer neighbor (pink) is pointing towards the reference CTC1 (cyan)-bound ssDNA, with the black dashed lines representing feasible ionic interactions between R1175 and phosphodiester groups of the ssDNA. **(C & D)** Identified inter-molecular interactions between CTC1, STN1 and TEN1 at interfaces of the decameric supercomplex.

A comparison of the monomeric DNA-free CST model (fig. S1) and the monomeric CST extracted from the decameric DNA-bound CST supercomplex (**Fig. 1D**) suggested that STN1c has two docking sites on CTC1. To investigate this, we returned to a cryo-EM dataset obtained during our buffer optimization phase for decameric CST that has a high population of monomeric CST with telomeric ssDNA added (fig. S12), and we performed further 3D classification on this monomeric DNA-bound CST. As expected, we found two distinct conformations – one with a “head” density and the other with an “arm” density, both ~25% of the original particle population – albeit at a lower model resolution of ~ 10 Å (**Fig. 6A & B**). Since STN1c is in the “arm” conformation in the decameric CST (**Fig. 5A**), and the STN1c “head” conformation would sterically hinder formation of the decamer (obstructing dihedral dimerization), we propose that switching from “head” to “arm” docking position for STN1c is an important first step for CST to form a decameric supercomplex. Surface-charge analysis revealed a highly positively charged surface on CTC1 OB-G, where the STN1n is expected to dock in the “head” conformation (**Fig. 6C**), and, interestingly, similar analysis revealed a reciprocal patch of highly negatively charged surface on STN1c (**Fig. 6C**, **inset**). This suggests charge-charge interactions could mediate the transition from monomeric to decameric CST, which would explain how a longer ssDNA, with extended binding to the OB-G’s negative-patch, can trigger this transition. The charge-charge interactions also suggest elevated salt could mediate the transition in the absence of ssDNA. Indeed, we saw a significant increase in decameric CST population without addition of ssDNA in non-physiological salt concentration of 800 mM NaCl (fig. S13).

**Fig. 6.**
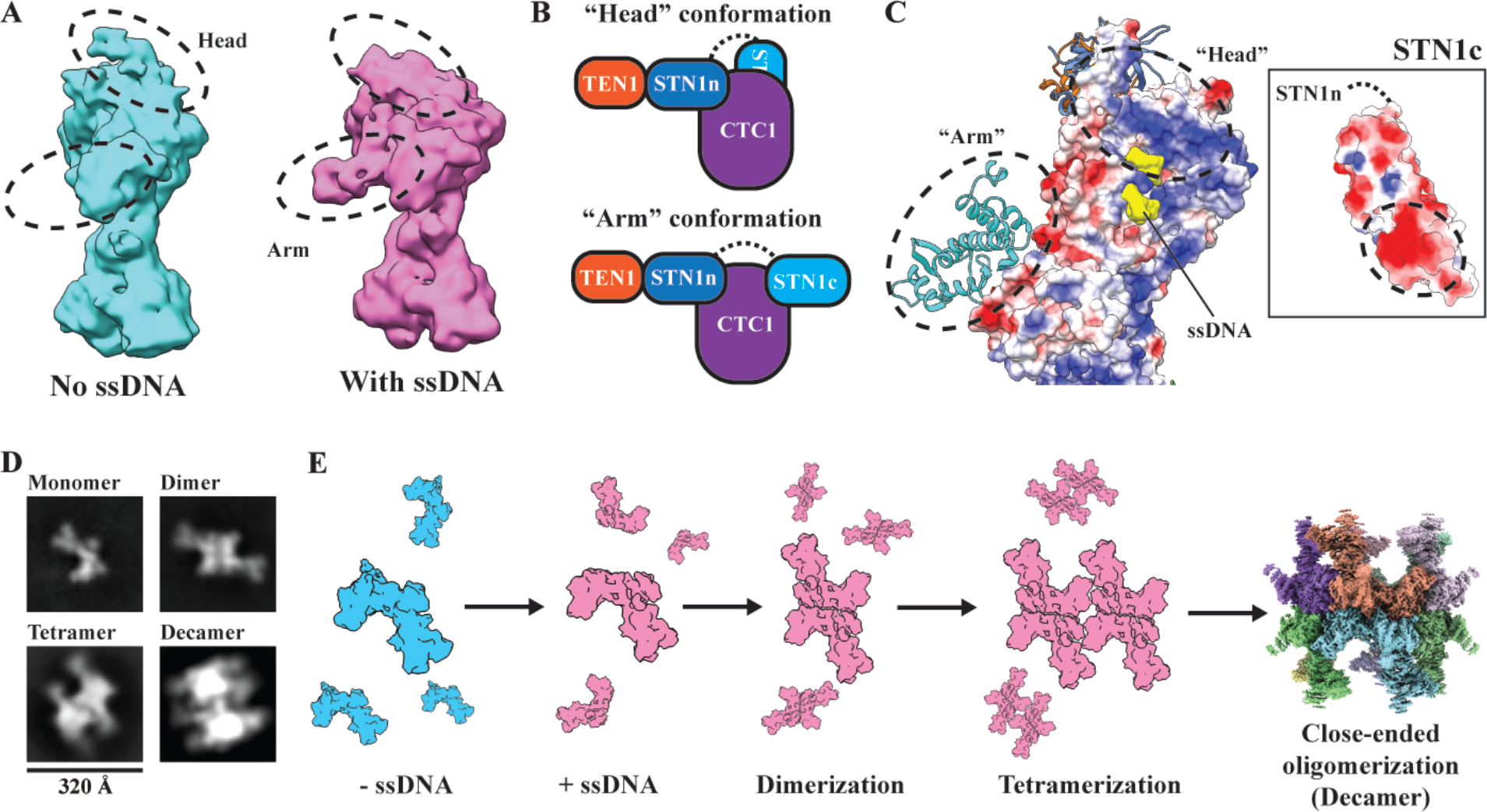
Assembly mechanism and pathway model of CST decameric supercomplex. **(A)** Cryo-EM densities of two conformations of monomeric CST with the differences indicated by dashed black circles. The two conformations – “head” (cyan) and “arm” (pink) – are assigned as CST without and with ssDNA bound, respectively. **(B)** Cartoon models of CST “head” and “arm” conformations depicted by conformational changes of STN1c docking site on CTC1. The black dashed line represents the unstructured polypeptide region between STN1n and STN1c. **(C)** Columbic surface analysis reveals a highly positively charged patch on CTC1 OB-G, where STN1c lies when in “head” conformation. Reciprocally, a highly negatively charged surface is shown on STN1c, see inset. **(D)** 2D classes of negative-stained CST incubated with 3xTEL ssDNA shows multiple oligomeric species of CST, which are assigned as monomer, dimer, tetramer and decamer. **(E)** Proposed model of assembly pathway of CST decameric supercomplex upon ssDNA introduction. CST binding of ssDNA prevents STN1c from binding to its original site (“head” conformation, cyan), allowing CST to form dimers before progressing to tetramers, and eventually leading to a close-ended decameric supercomplex.

Finally, we used negative-stain EM single-particle analysis to identify subcomplexes of the decamer that would give hints to its assembly pathway(s). We observed two homogenous subcomplexes; dimers and tetramers (**Fig. 6D**), which are plausible intermediates in an assembly pathway such as the following: CST assembles first via a head-head dimer (dihedral dimer) before forming a lateral tetramer involving two dihedral dimers, and eventually closing the symmetric circle by continuing the lateral oligomerization till a decameric supercomplex is formed (**Fig. 6D & E**). This assembly of CST into a decameric supercomplex is unexpected, but, intriguingly, reminiscent of the oligomeric nature of some DNA replication machineries (*42*, *43*). Given the clustering nature of DNA damage repair centers and replication forks (*44*, *45*), a key future research direction will be to determine the role(s) that CST decameric formation may play in functional molecular aggregation.

## Discussion

Our structure of the decameric human CST supercomplex bound with telomeric ssDNA provides the platform for understanding mechanisms of various CST functions in DNA replication and DNA damage repair, not only at telomeres but also genome-wide (*9*, *10*, *13*, *15*, *46*). This work showcases an unprecedented assembly pathway where a two-megadalton protein supercomplex is triggered by DNA binding, and it further demonstrates the utility of cryo-EM single-particle analysis for determining the atomic model of a protein structure as large as ~200 kDa. The atomic-resolution details revealed in this structure have enabled us to identify amino acids responsible for the numerous interactions that are key for CST functions, i.e., the intricate heterotrimer assembly, ssDNA-binding anchor site and decamer assembly. With genetic engineering, we and others can now systematically mutate these residues in cells/organisms and study their phenotypes, especially as the molecular function(s) of the decameric form of CST is still unclear. It will also serve as a molecular model to understand the underlying mechanisms of CST-related human diseases such as Coats Plus and Dyskeratosis congenita.

Our work also shows Cdc13 is poorly homologous to CTC1 not only in terms of sequence but also structurally. CTC1 is more similar to a duplicated form of RPA70, providing insights to CST evolutionary pathways. The decameric form of CST with ssDNA binding capacity of up to ten telomeric repeats of ssDNA suggests a tantalizing possibility of CST organizing telomere overhangs into compact and restrictive structures, similar to nucleosomal organization of dsDNA. This aligns well with CST end-protection functions at telomeres and sequestration of ssDNA at stalled replication forks genome-wide (*47*).

## Acknowledgments

We thank Z. Yu, D. Matthies, and R. Huang (Janelia Research Campus Cryo-EM facility), P. Blerkom and J. Kieft (University of Colorado Anschutz EM facility) and G. Morgan and C. Page (University of Colorado Boulder EM facility) for help in microscope setup and data collection. We thank F. Asturias and T. Terwilliger for stimulating discussions. We thank T. Nahreini (University of Colorado Boulder, Department of Biochemistry Cell Culture Facility) and BioFrontiers Institute Computing Core for their support and assistance in this work. We thank the Cech lab and Wuttke lab members for their helpful suggestions.

## Funding

This work is funded in part by grants from National Institutes of Health (R01GM099705) to T.R.C, (R01GM059414) to D.S.W, (K99GM131023) to C.L., and the National Science Foundation (MCB 1716425) to D.S.W. A.T.B. is supported by a fellowship provided by National Institutes of Health/University of Colorado Boulder, Signaling and Cellular Regulation Graduate Training grant (T32GM008759). T.R.C is a Howard Hughes Medical Institute Investigator.

## Author Contributions

C.L., D.S.W and T.R.C. outlined the experimental plans. C.L. established experimental procedures, expressed and purified the complexes from insect cells, optimized conditions for cryo-EM work, prepared cryo-EM grids, collected and processed EM datasets, did the model building, refinement and validations. A.T.B. assisted in insect cell expression and purification, negative-stain screening and analysis, and did gel-shifts assays. A.J.Z., A.E.M and C.L. performed molecular cloning, expressed and purified complexes in human cultured cells for pull-down experiments and analysis. C.L., D.S.W. and T.R.C. analyzed and interpreted the model. C.L. and T.R.C wrote the manuscript with input from all authors.

## Competing interests

T.R.C. is on the board of directors of Merck and a consultant for STORM Therapeutics.

