## Supplementary Materials for "The cryo-EM structure of human CST reveals a two-megadalton decameric assembly bound to telomeric DNA"

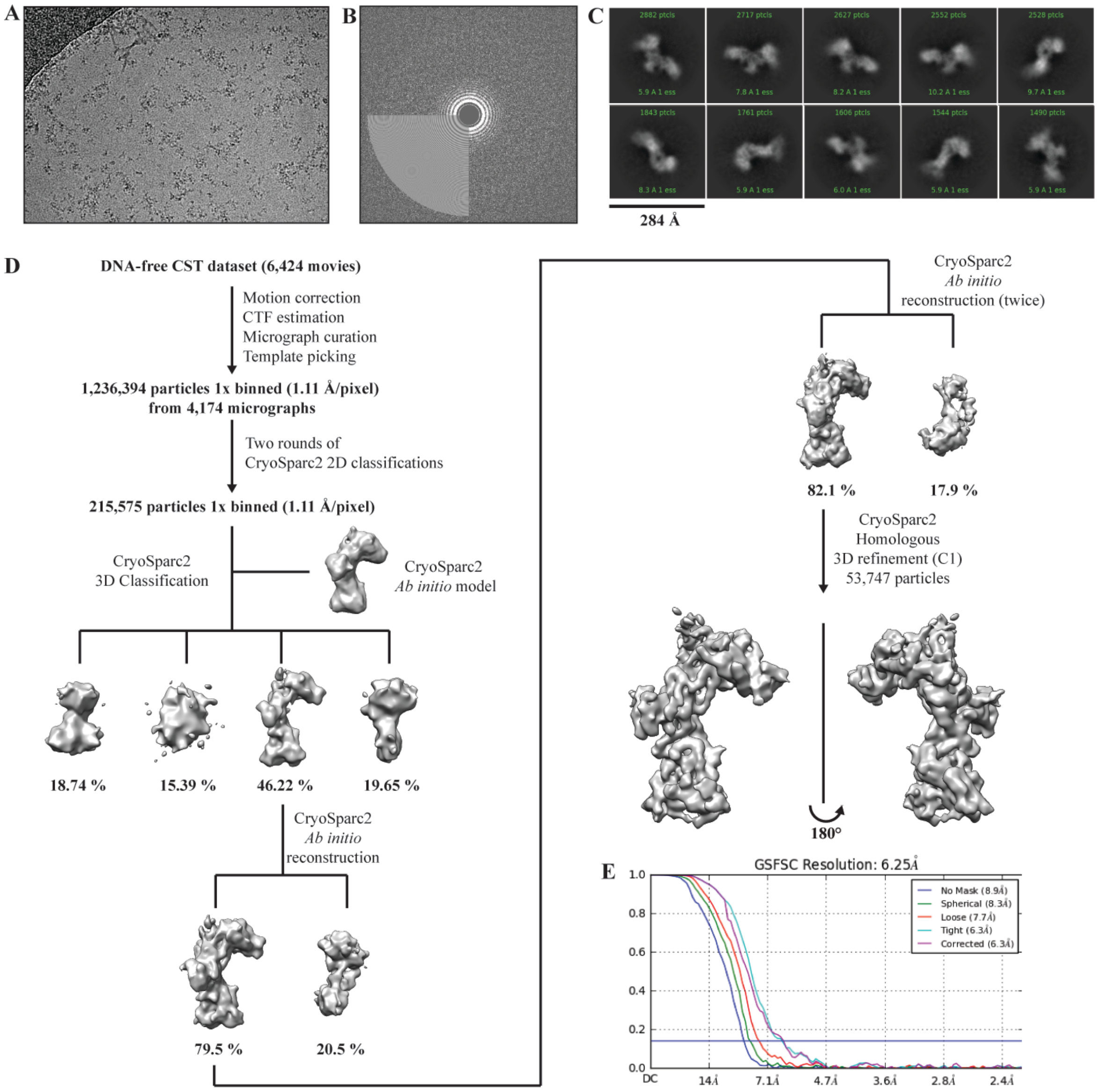


**Fig. S1. Cryo-EM reconstruction of DNA-free monomeric CST complex. (A)** A representative motion corrected micrograph (denoised by JANNI (61)). **(B)** A representative calculated CTF image. **(C)** Top ten 2D classes selected after the final 2D classification step, showing various orientations of DNA-free CST complex. **(D)** Cryo-EM processing pipeline of DNA-free CST. **(E)** CryoSparc2 (56) corrected Fourier shell correlation (FSC) reports a global resolution of 6.25 Å (purple curve).

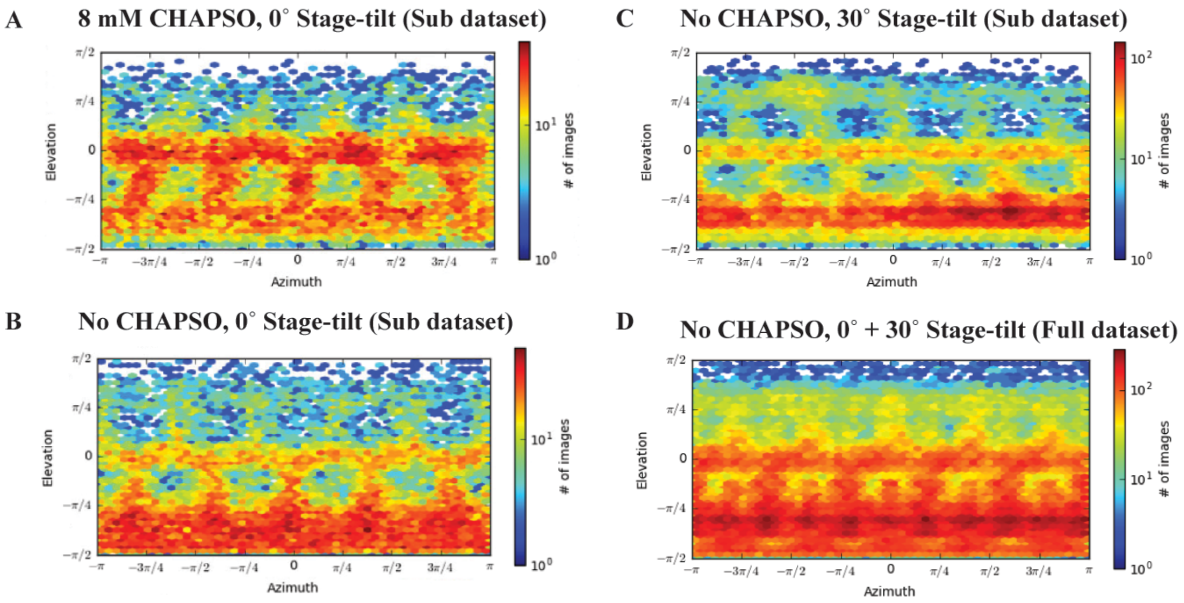


**Fig. S2. Viewing distributions of CST-3xTEL decamer in presence of CHAPSO or stage-tilt.** Distributions of particles orientations from 3D refinement step were used as an analysis to determine optimal data collection strategy. Panels A-C were calculated from sub datasets (~50,000 particles each) for fair comparison. In the presence of 8 mM CHAPSO **(A)**, the decamer orientation was more evenly distributed than that without CHAPSO added **(B & C)** but also caused particles aggregation, reducing data collection efficiency. To circumvent preferred orientations issue without utilizing CHAPSO, we adopted a two-pronged approach by collecting CST-3xTEL decamer at 30° stage-tilt (see top right panel;) and increasing the particle count (brute force) at 0° stage-tilt (> 20,000 movies). The panel D shows the final viewing distribution (D5 symmetry refinement) of the combined 0° + 30° stage-tilt full datasets (~ 300,000 particles). All figures were generated from CryoSparc2 (56).

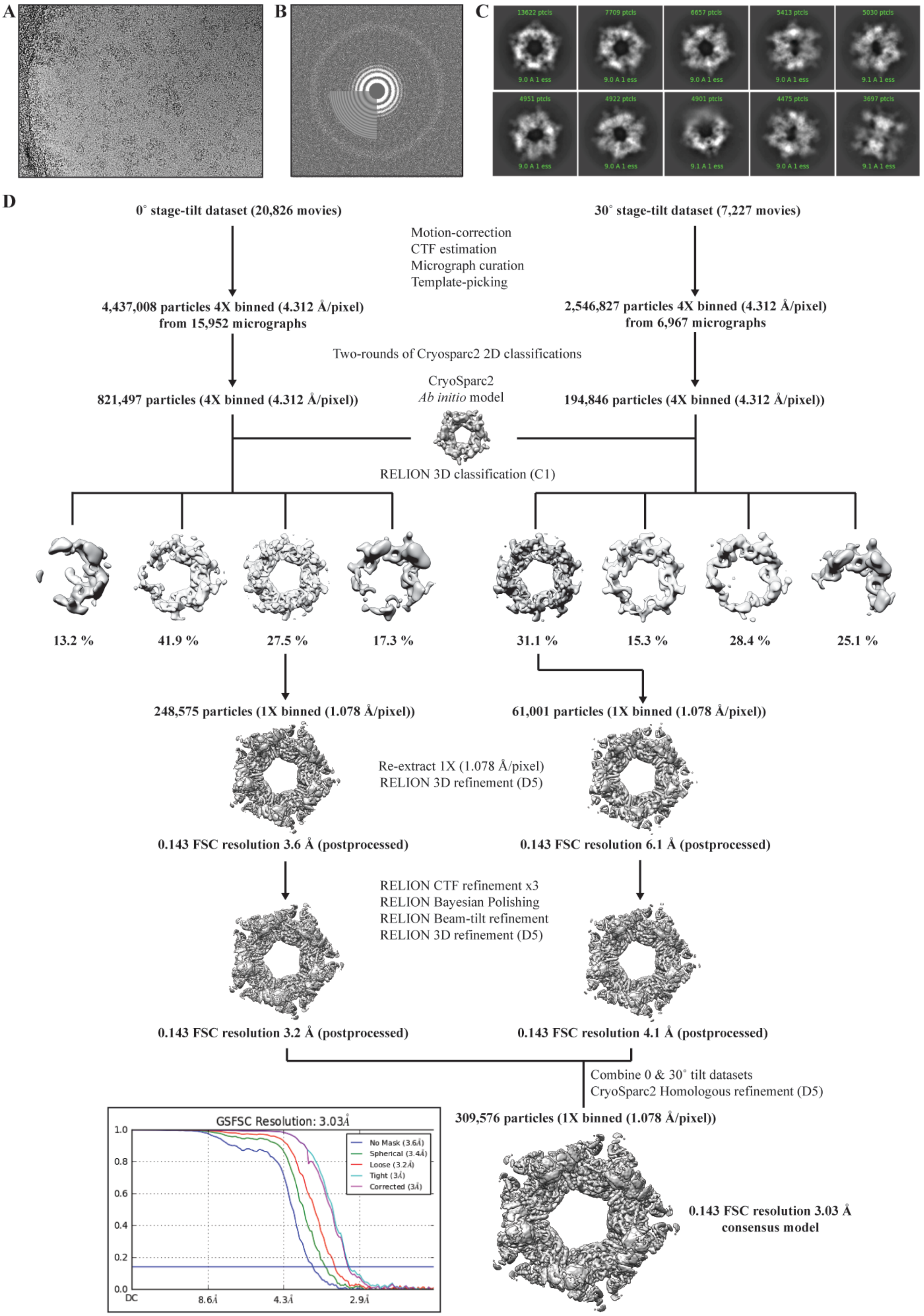
A
B
C
13622 ptcls
7709 ptcls
6657 ptcls
5413 ptcls
5030 ptcls
4951 ptcls
4922 ptcls
4901 ptcls
4475 ptcls
3697 ptcls
D
0˚ stage-tilt dataset (20,826 movies)
30˚ stage-tilt dataset (7,227 movies)
Motion-correction
CTF estimation
Micrograph curation
Template-picking
4,437,008 particles 4X binned (4.312 Å/pixel) from 15,952 micrographs
2,546,827 particles 4X binned (4.312 Å/pixel) from 6,967 micrographs
Two-rounds of Cryosparc2 2D classifications
821,497 particles (4X binned (4.312 Å/pixel))
CryoSparc2
Ab initio model
194,846 particles (4X binned (4.312 Å/pixel))
RELION 3D classification (C1)
13.2 %
41.9 %
27.5 %
17.3 %
31.1 %
15.3 %
28.4 %
25.1 %
248,575 particles (1X binned (1.078 Å/pixel))
61,001 particles (1X binned (1.078 Å/pixel))
Re-extract 1X (1.078 Å/pixel)
RELION 3D refinement (D5)
0.143 FSC resolution 3.6 Å (postprocessed)
0.143 FSC resolution 6.1 Å (postprocessed)
RELION CTF refinement x3
RELION Bayesian Polishing
RELION Beam-tilt refinement
RELION 3D refinement (D5)
0.143 FSC resolution 3.2 Å (postprocessed)
0.143 FSC resolution 4.1 Å (postprocessed)
Combine 0 & 30˚ tilt datasets
CryoSparc2 Homologous refinement (D5)
309,576 particles (1X binned (1.078 Å/pixel))
GSFSC Resolution: 3.03Å
No Mask (3.6Å)
Spherical (3.4Å)
Loose (3.2Å)
Tight (3Å)
Corrected (3Å)
1.0
0.8
0.6
0.4
0.2
0.0
DC
8.6Å
4.3Å
2.9Å
0.143 FSC resolution 3.03 Å consensus model

**Fig. S3. Cryo-EM reconstruction of CST-3xTEL decamer complex. (A)** A representative motion corrected micrograph. **(B)** A representative calculated CTF image with water diffraction ring visible. **(C)** Top ten 2D classes selected after the final 2D classification step, showing various orientations of CST-3xTEL decamer complex. **(D)** Cryo-EM processing pipeline of stage-tilt datasets for CST-3xTEL decamer samples. Inset shows CryoSparc2 (56) corrected Fourier shell correlation (FSC) plot which reports a global resolution of 3.03 Å (purple curve).

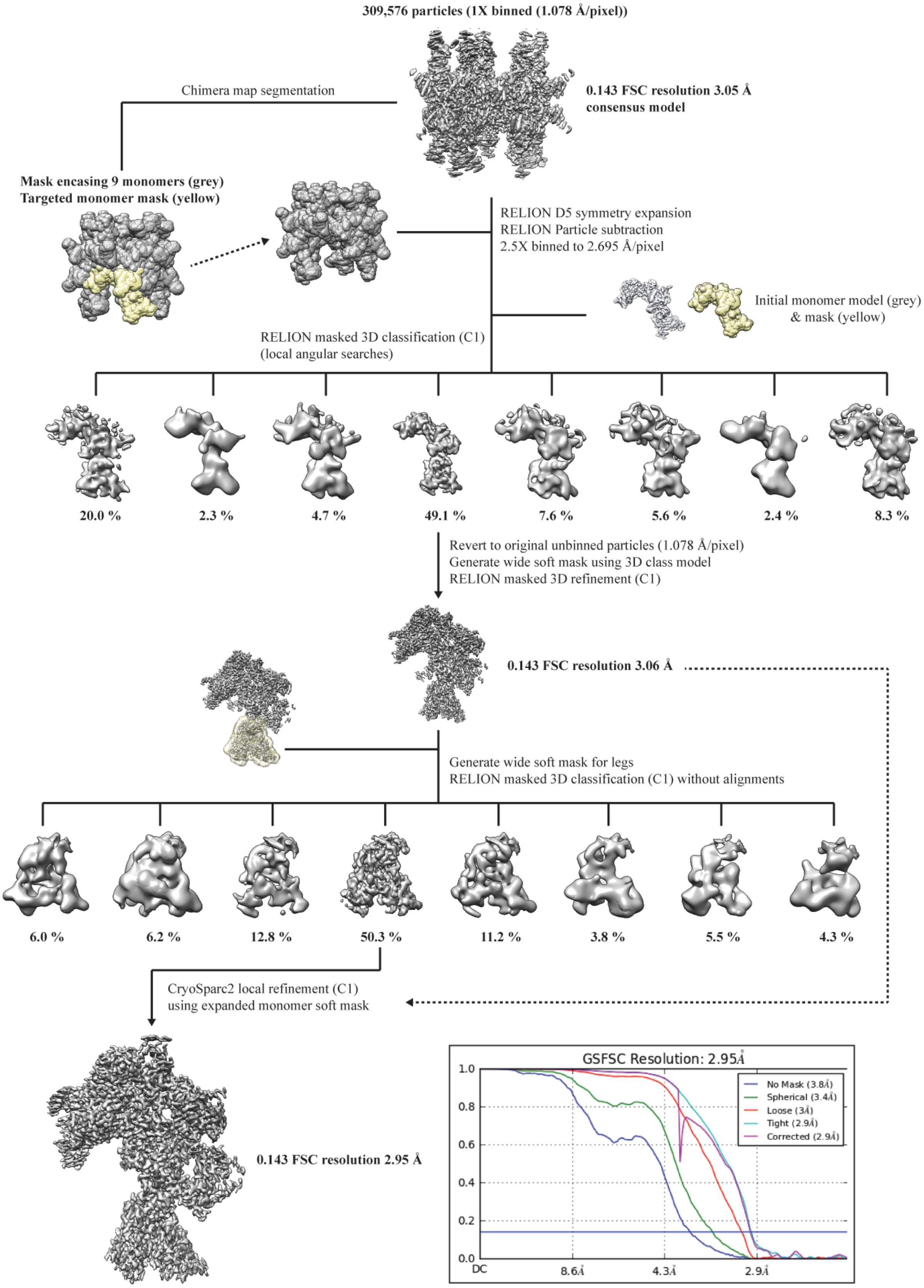
309,576 particles (1X binned (1.078 Å/pixel))
0.143 FSC resolution 3.05 Å
consensus model
Chimera map segmentation
Mask encasing 9 monomers (grey)
Targeted monomer mask (yellow)
RELION D5 symmetry expansion
RELION Particle subtraction
2.5X binned to 2.695 Å/pixel
Initial monomer model (grey)
& mask (yellow)
RELION masked 3D classification (C1)
(local angular searches)
20.0 %
2.3 %
4.7 %
49.1 %
7.6 %
5.6 %
2.4 %
8.3 %
Revert to original unbinned particles (1.078 Å/pixel)
Generate wide soft mask using 3D class model
RELION masked 3D refinement (C1)
0.143 FSC resolution 3.06 Å
Generate wide soft mask for legs
RELION masked 3D classification (C1) without alignments
6.0 %
6.2 %
12.8 %
50.3 %
11.2 %
3.8 %
5.5 %
4.3 %
CryoSparc2 local refinement (C1)
using expanded monomer soft mask
0.143 FSC resolution 2.95 Å
GSFSC Resolution: 2.95Å
No Mask (3.8Å)
Spherical (3.4Å)
Loose (3Å)
Tight (2.9Å)
Corrected (2.9Å)
1.0
0.8
0.6
0.4
0.2
0.0
DC
8.6Å
4.3Å
2.9Å

**Figure S4. Conformational extraction of CST monomer from decamer complex for cryo-EM reconstruction.** The DNA-free CST monomer model (Fig. S1) was used as a guide to segment the decamer mask and remove a single monomer, resulting in a "10-1" soft mask. This mask was applied to the D5 symmetry expanded particle stack for particle subtraction, effectively removing nine CST monomers and leaving one monomer in a standardized position in the subtracted particle stack. 3D classification was used to identify a homogenous conformation for a CST monomer, which was carried forward for 3D refinement, leading to a 3.06 Å map. A soft mask for the legs (see main text for description) was created to focus on isolating a population of particles with well-resolved legs region by 3D classification without alignments. The particles subset from a single 3D class showing well-defined "legs" were used for 3D refinement in CryoSparc2, leading to the final 2.95 Å map used for *de novo* building.

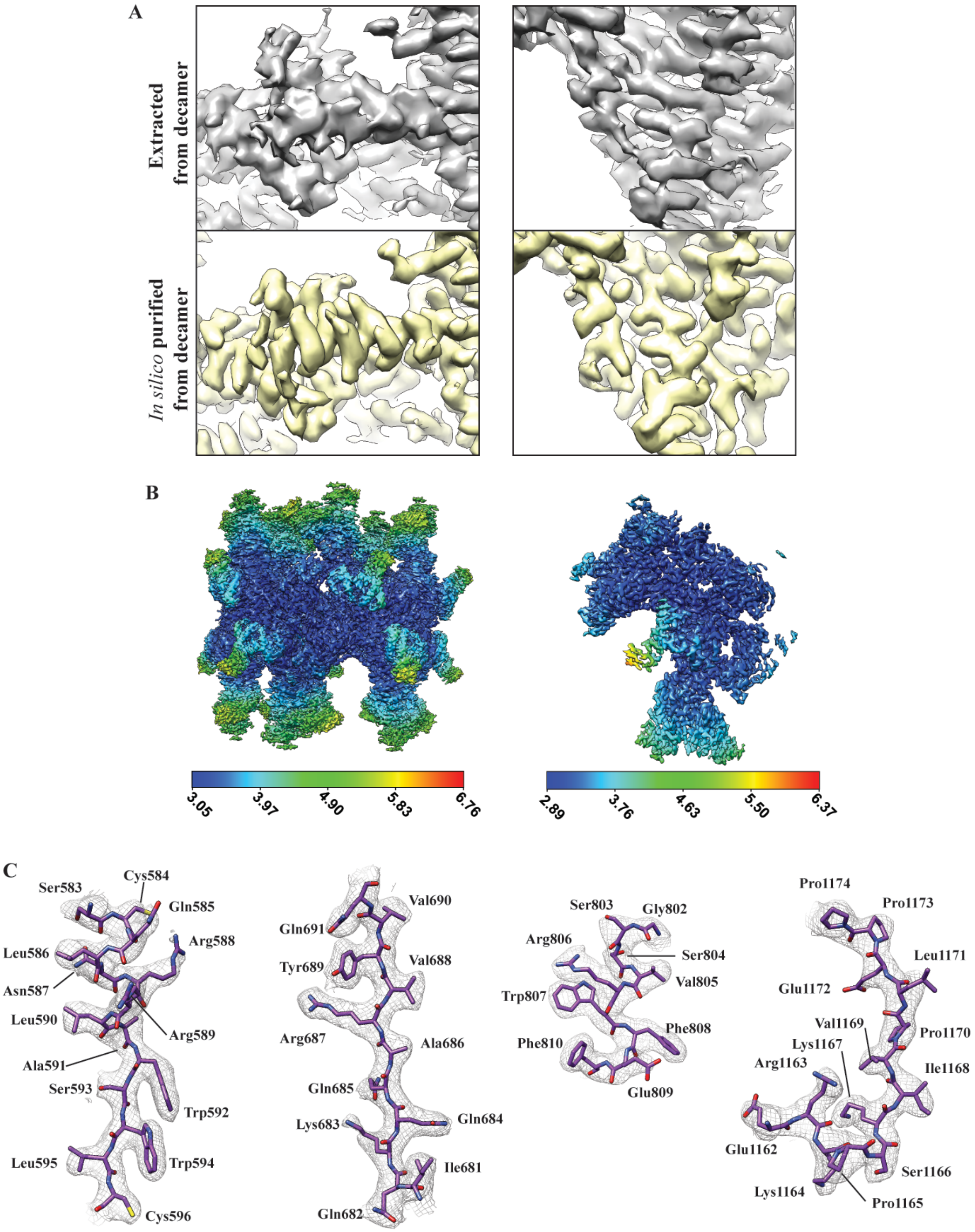


**Figure S5. Quality differences between cryo-EM maps after *in silico* purification of CST monomer from decamer map and *de novo* model building. (A)** Comparison of the cryo-EM maps of CST monomer extracted directly from decamer consensus map (grey density) and that *in silico* purified (yellow density, see Fig. S4 for details) showing significant improvements in map interpretation. **(B)** Local resolution mapping of CST-3xTEL decamer consensus model (left panel) and after *in silico* purification (right panel). **(C)** Representative EM map density encasing the *de novo* built atomic models of CTC1.

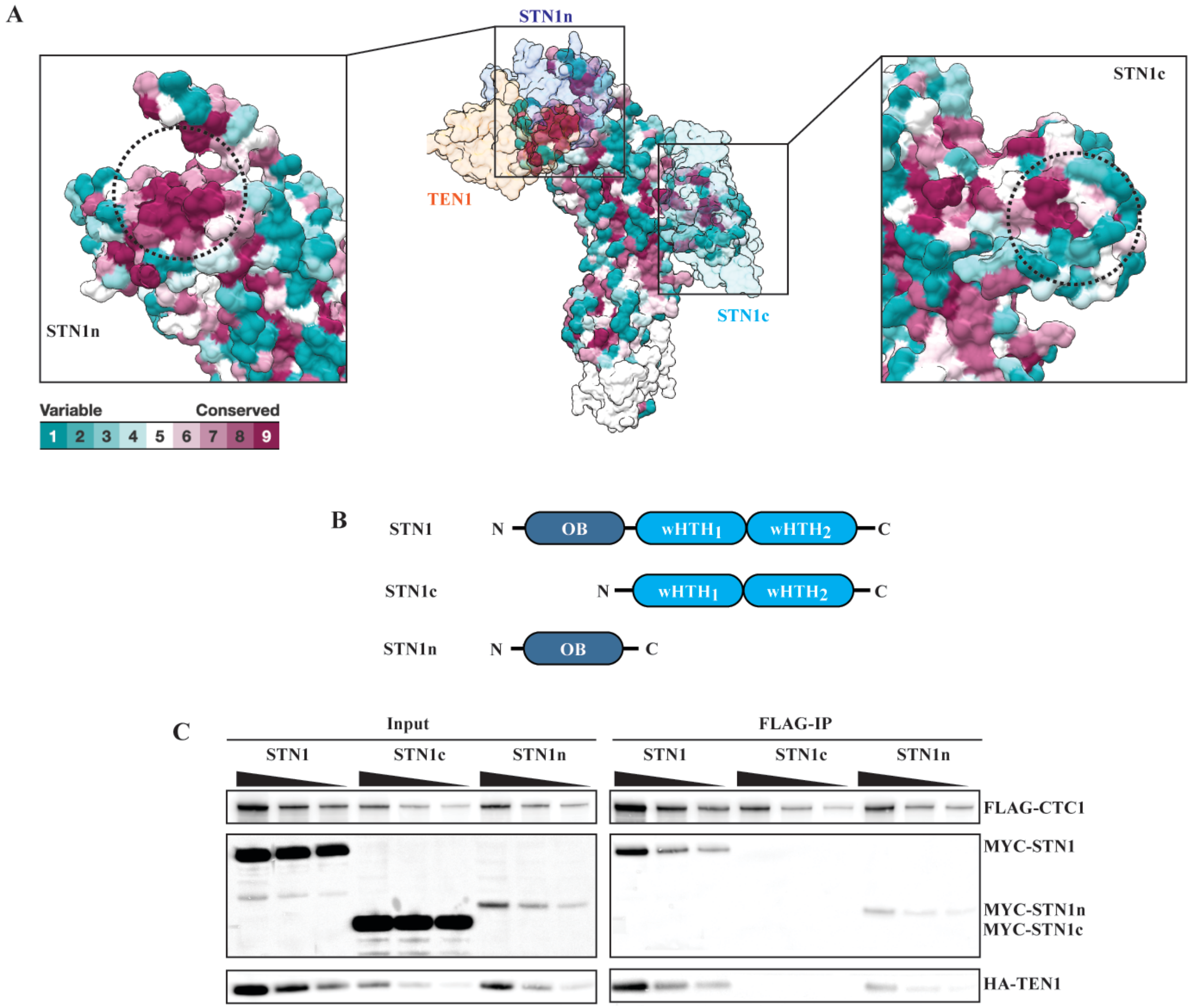


**Figure S6. STN1 anchor site on CTC1. (A & B)** ConSurf (72) analysis showing the conservation of CTC1 residues at the interaction sites of STN1 two subdomains – STN1c and STN1n. **(C)** FLAG-immunoprecipitation (FLAG-IP) of full-length STN1, STN1c and STN1n with 3xFLAG-CTC1 co-expressed in HEK293T cells. From input samples, STN1n was less stably expressed than full-length STN1 and STN1c but was still able to bind CTC1 and TEN1, while STN1c did not stably associate.

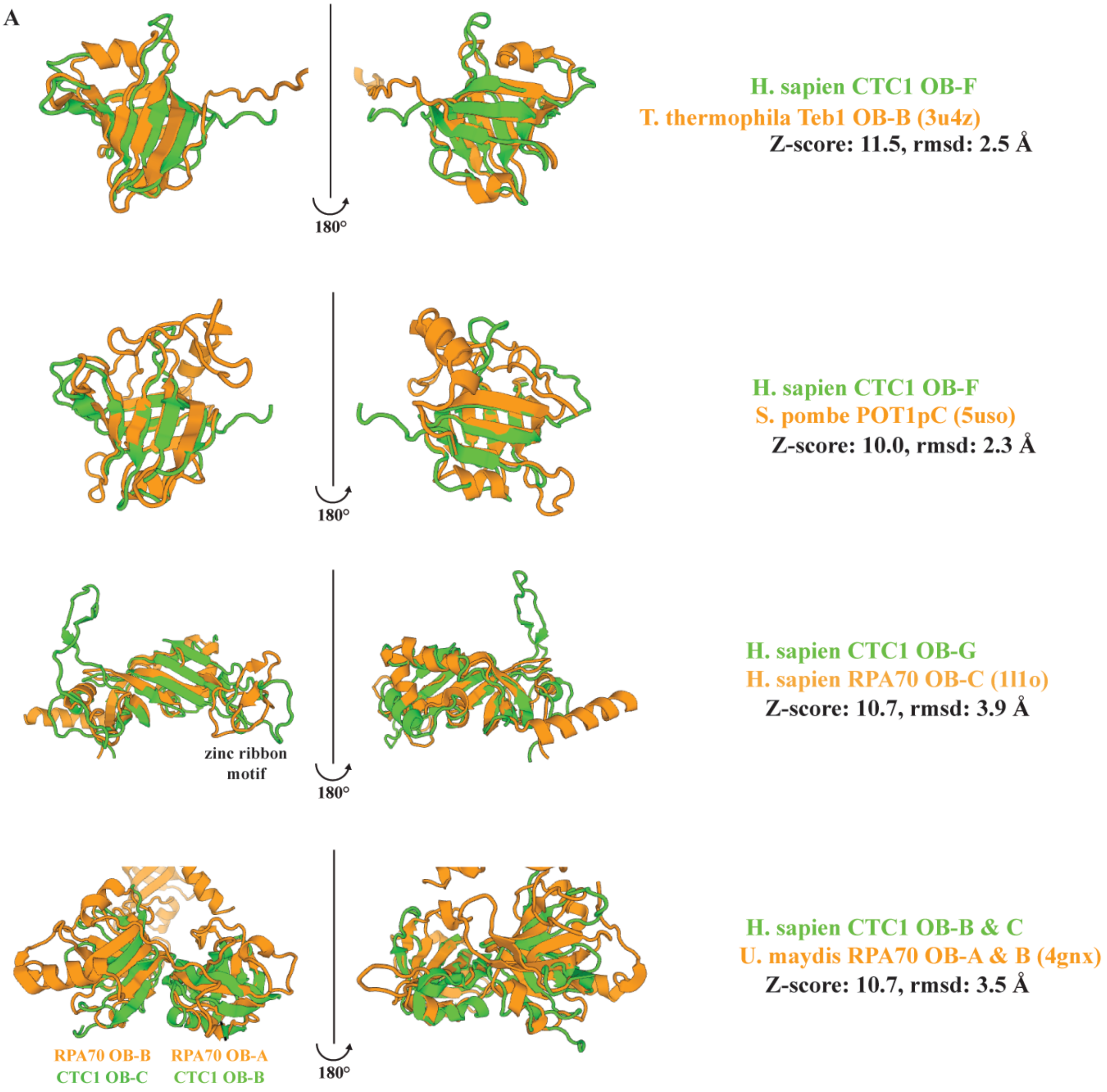
A
180°
H. sapien CTC1 OB-F
T. thermophila Teb1 OB-B (3u4z)
Z-score: 11.5, rmsd: 2.5 Å
H. sapien CTC1 OB-F
S. pombe POT1pC (5uso)
Z-score: 10.0, rmsd: 2.3 Å
zinc ribbon motif
H. sapien CTC1 OB-G
H. sapien RPA70 OB-C (1l1o)
Z-score: 10.7, rmsd: 3.9 Å
RPA70 OB-B
CTC1 OB-C
RPA70 OB-A
CTC1 OB-B
H. sapien CTC1 OB-B & C
U. maydis RPA70 OB-A & B (4gnx)
Z-score: 10.7, rmsd: 3.5 Å

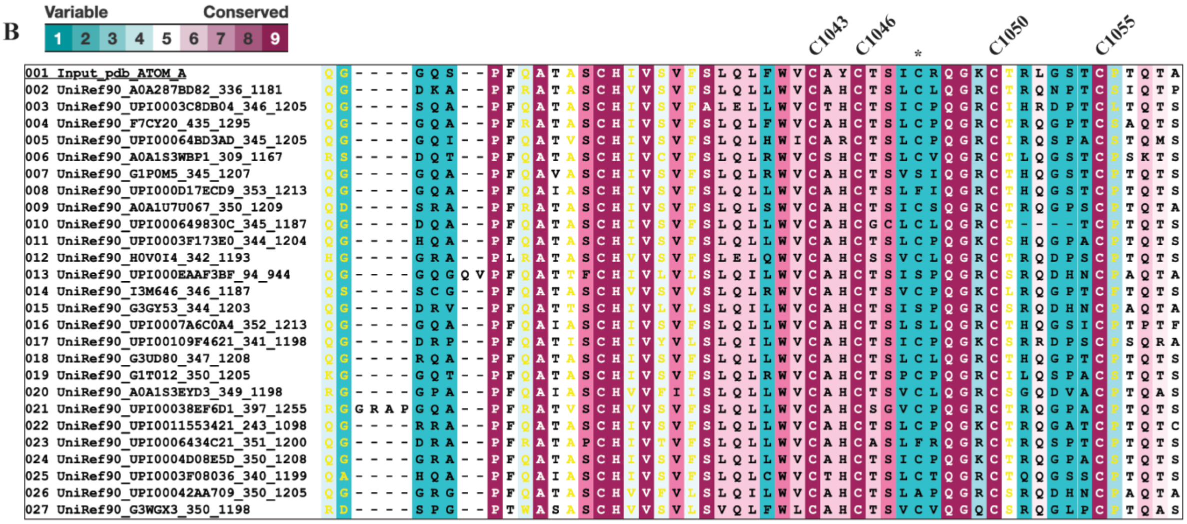
B
Variable
Conserved
1 2 3 4 5 6 7 8 9
C1043
C1046
C1050
C1055

**Figure S7. Structural homology search and conservation of CTC1 zinc ribbon motif. (A)** Top relevant structural homology hits of CTC1 OB-domains (B, C, F & G). DALI (71) was used to search for structural homology. **(B)** Cysteines involved in zinc coordination in the zinc ribbon motif are highly conserved. ConSurf (72) was used for conservation analysis.

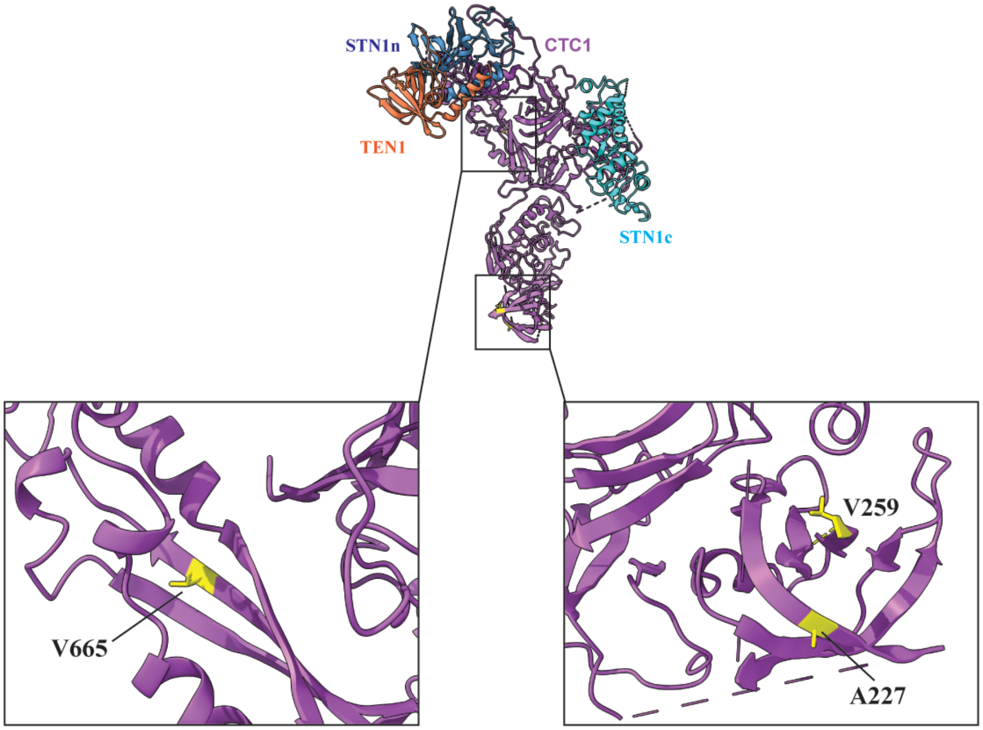


**Figure S8. Structural mapping of CTC1 disease mutations that affect CST pol-α interaction.** CTC1 residues (yellow) that affect pol-α interaction with CST (A227, V259, V665) (6) are located in two separate sites.

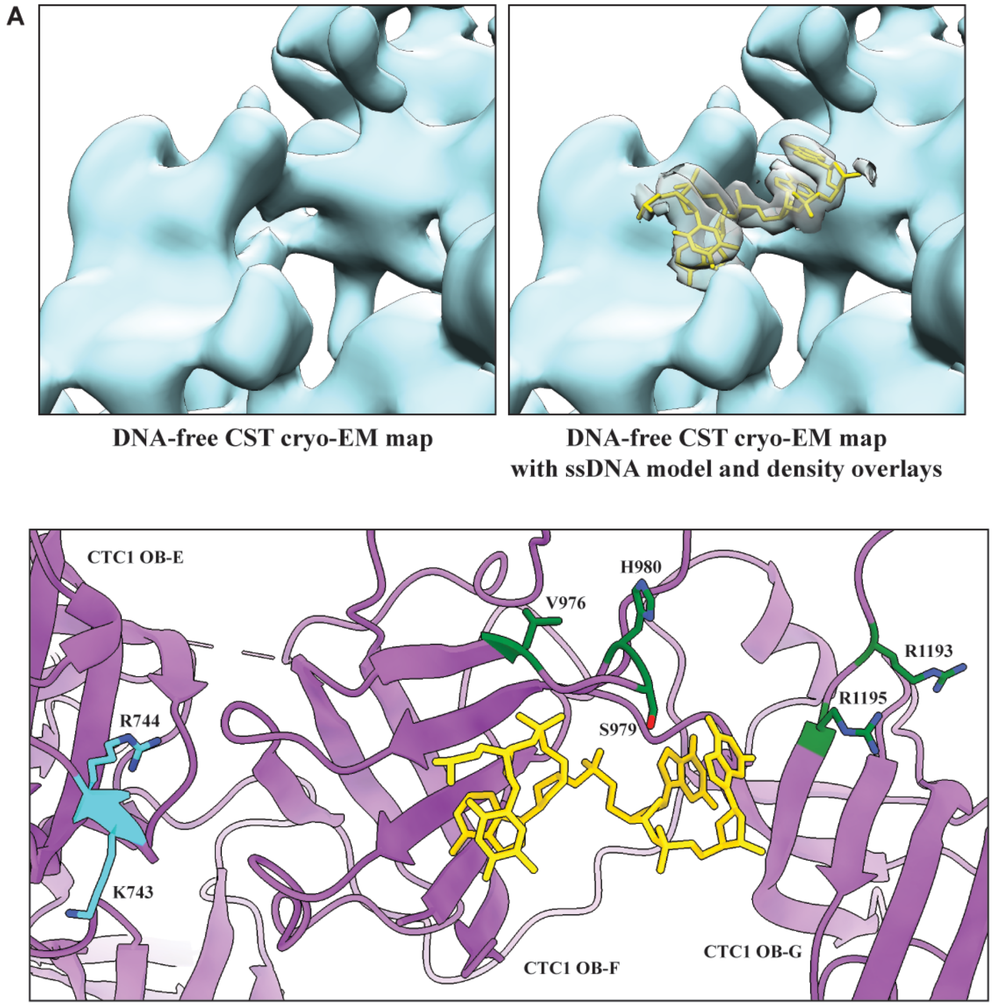


**Figure S9. Single-stranded DNA EM density was not seen in DNA-free CST EM map and structural mapping of additional CTC1 residues chosen for DNA-binding mutagenesis. (A)** The DNA-free CST EM map (turquoise, left panel) did not have the extra density of ssDNA (grey with yellow atomic model of ssDNA, right panel) that was found in the map of CST-3xTEL decamer. **(B)** Additional CTC1 amino acid residues (green or cyan) chosen for DNA-binding mutagenesis. Green residues were DNA-binding defective upon mutagenesis. Cyan residues selected for DNA-binding control (see main text).

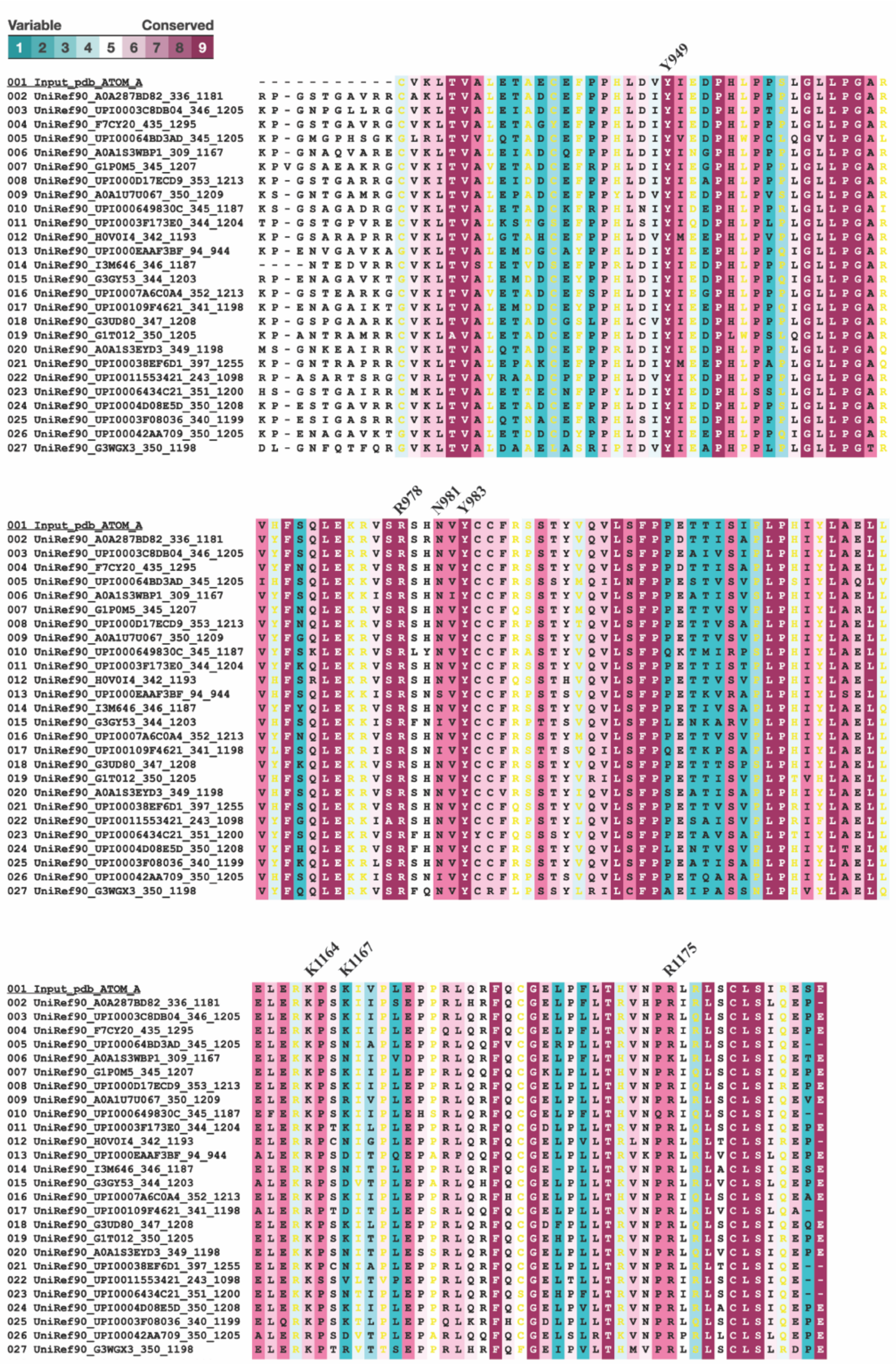

**Figure S10. Residues conservation analysis of CTC1 DNA-binding mutants.** ConSurf (72) was used to determine the conservation of identified DNA-binding residues of CTC1. Although CTC1 K1167 apparently has poor residue conservation, the variations were limited to mostly charged residues.

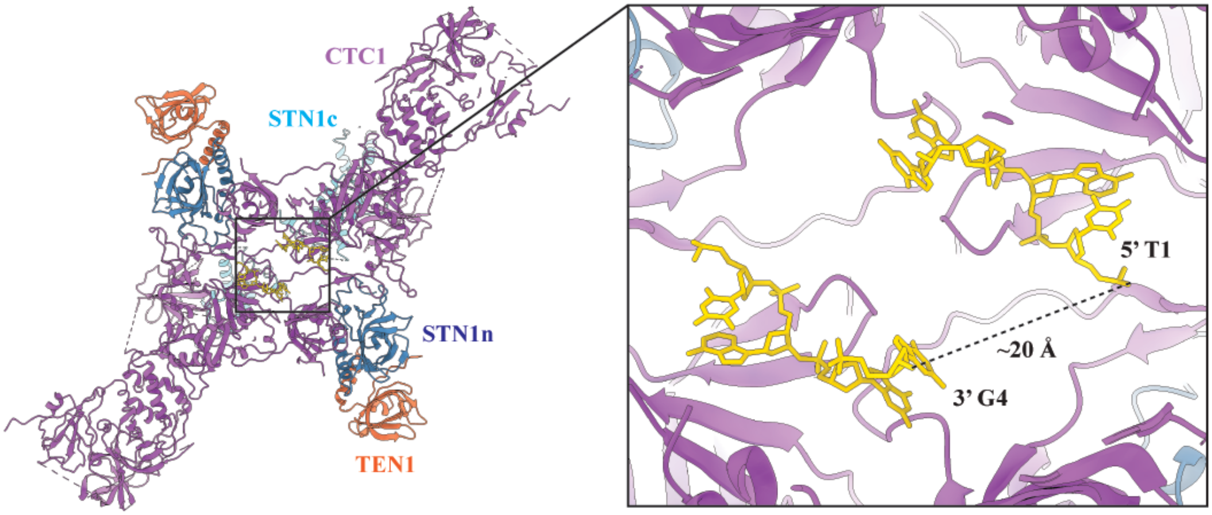


**Figure S11. Molecular distance between dihedral dimers' single-stranded DNA molecules.** Left panel shows a CST dihedral dimer (see main text for details). Right panel is the zoomed-in image of the two single-stranded DNA (ssDNA) molecules belonging to the dimer. Measured distance between the 5' T1 and 3' G4 of the opposite ssDNA (yellow) is approximately 20 Å. A length of 20 Å will be equivalent to approximately 3 nucleotides (assuming a nucleotide is 7 Å long). This is sufficient for a single $(TTAGGG)_3$ ssDNA molecule to engage both DNA anchor sites of a CST dihedral dimer.

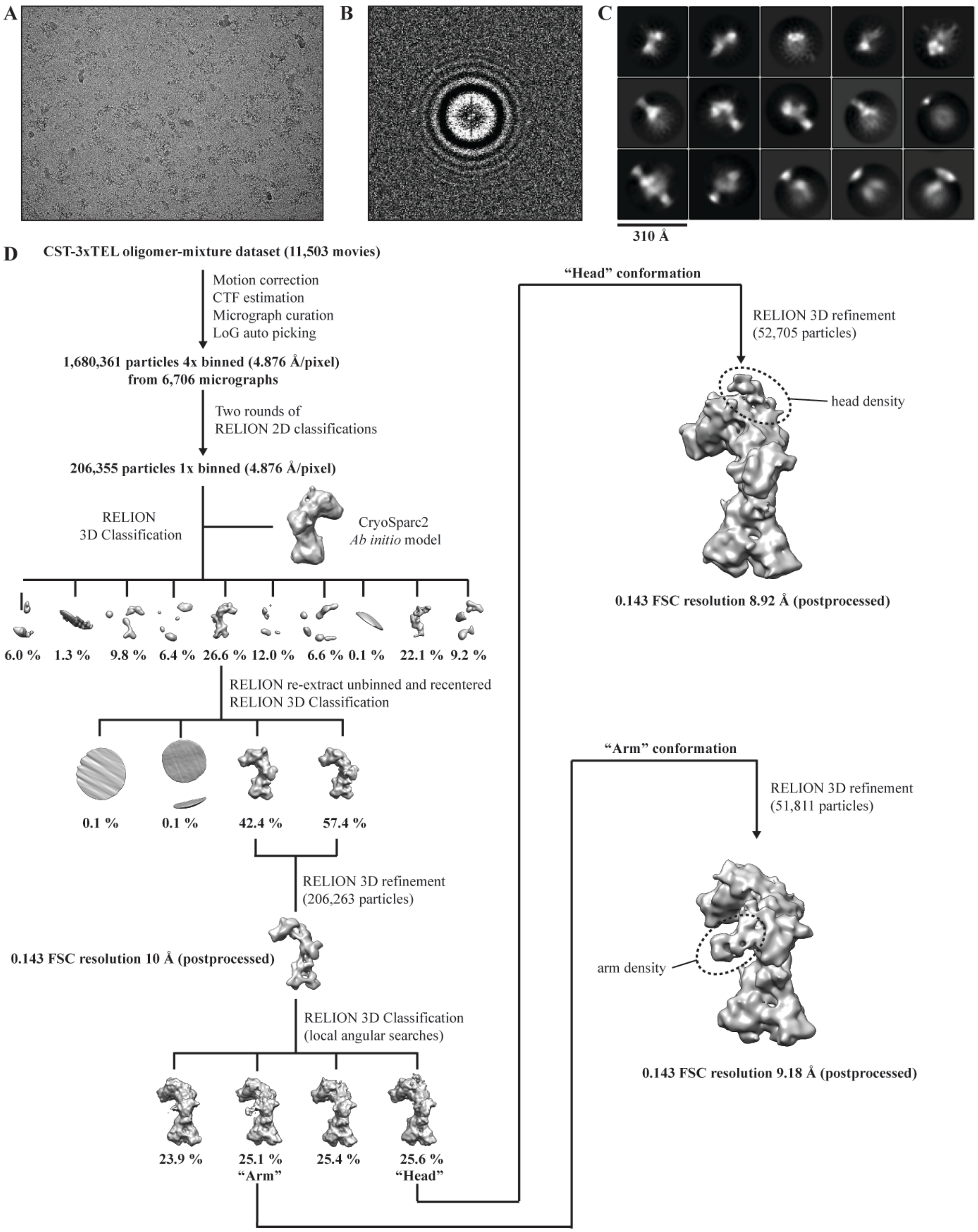


**Fig. S12. Cryo-EM reconstruction of CST-3xTEL sample showing two distinct conformations. (A)** A representative motion-corrected micrograph. **(B)** A representative calculated CTF image. **(C)** Top fifteen 2D classes selected after the final 2D classification step. **(D)** Cryo-EM processing pipeline of DNA-free CST.

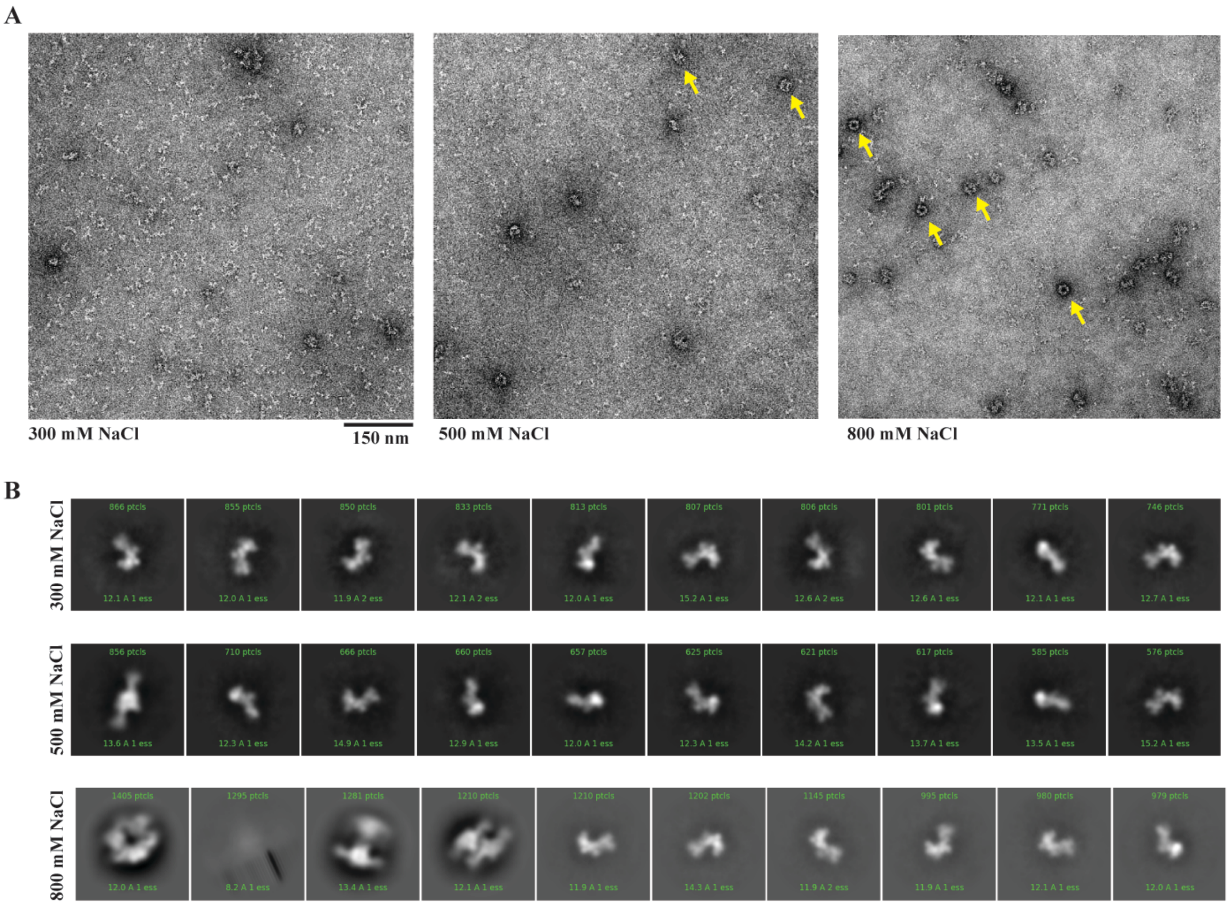


**Figure S13. Negative-stain EM analysis of CST decamer formation salt-dependence. (A)** Representative micrographs of CST incubated in buffers containing 300, 500 and 800 mM NaCl salt. Yellow arrows indicate CST decamer particles. **(B)** 2D classifications showing the top ten classes of particles (based on particles distribution). CST starts to form decameric particles with increasing NaCl concentration in buffer.

| CST complexes | DNA-free CST | CST-3xTEL oligomer-mixture | CST-3xTEL Decamer (0° tilt) | CST-3xTEL Decamer (30° tilt) | CST-3xTEL Decamer (0° + 30° tilt + D5 symmetry expanded) |
|---|---|---|---|---|---|
| **Data collection and processing** | | | | | |
| Magnification | 36,000 | 29,000 | 81,000 | 81,000 | 81,000 |
| Voltage (kV) | 200 | 200 | 300 | 300 | 300 |
| Electron exposure (e–/Å$^2$) | 53.6 | 55.0 | 60.0 | 60.0 | 60.0 |
| Defocus range (μm) | -2.0 to -3.5 | -1.5 to -3.5 | -1.0 to -2.5 | -1.0 to -2.5 | -1.0 to -2.5 |
| Pixel size (Å) | 1.11 (Count.) | 0.6095 (Super.) | 0.539 (Super.) | 0.539 (Super.) | 0.539 (Super.) |
| Symmetry imposed | C1 | C1 | D5 | D5 | C1 |
| Initial particle images (no.) | 1,236,394 | 1,680,361 | 4,437,008 | 2,546,827 | 3,095,760 |
| Final particle images (no.) | 53,747 | 52,705, 51,811 | 248,575 | 61,001 | 833,627 |
| Map resolution (Å) | 6.25 | 8.92, 9.18 | 3.2 | 4.1 | 2.95 |
| FSC threshold | 0.143 | 0.143 | 0.143 | 0.143 | 0.143 |
| **Refinement** | | | | | |
| Initial model used (PDB code) | | | | | 5W2L, 4JOI, 4JQF |
| Model resolution (Å) | | | | | 3.30 |
| FSC threshold | | | | | 0.5 |
| Map sharpening B factor (Å$^2$) | | | | | -80 |
| Model composition | | | | | |
| Non-hydrogen atoms | | | | | 11,054 |
| Protein residues | | | | | 1397 |
| Ligands (DNA) | | | | | 4 |
| B factors (Å$^2$) | | | | | |
| Protein | | | | | 120.23 |
| Ligand (DNA) | | | | | 131.46 |
| R.m.s. deviations | | | | | |
| Bond lengths (Å) | | | | | 0.007 |
| Bond angles (°) | | | | | 1.037 |
| **Validation** | | | | | |
| MolProbity score | | | | | 1.89 |
| Clashscore | | | | | 5.24 |
| Poor rotamers (%) | | | | | 0.67 |
| Ramachandran plot | | | | | |
| Favored (%) | | | | | 87.62 |
| Allowed (%) | | | | | 12.16 |
| Disallowed (%) | | | | | 0.22 |
| EMRinger | | | | | 2.18 |

**Table S1. Cryo-EM data collection, refinement and validation statistics.**
